# Tissue specific LRRK2 interactomes reveal a distinct functional unit within the striatum

**DOI:** 10.1101/2022.06.28.497918

**Authors:** Yibo Zhao, Nikoleta Vavouraki, Ruth C Lovering, Valentina Escott-Price, Kirsten Harvey, Patrick A Lewis, Claudia Manzoni

## Abstract

Mutations in *LRRK2* are the most common genetic cause of Parkinson’s disease. Despite substantial research efforts, the physiological and pathological role of this multidomain protein remains poorly defined. In this study, we used a systematic approach to construct the general protein-protein interactome around LRRK2, which was then differentiated into 15 tissue-specific interactomes taking into consideration the differential expression patterns and the co-expression behaviours of the LRRK2 interactors in different healthy tissues. The LRRK2 interactors exhibited distinct expression features in the brain as compared to the peripheral tissues analysed. Moreover, a high degree of similarity was found for the LRRK2 interactors in *putamen, caudate* and *nucleus accumbens*, thus defining a potential LRRK2 functional cluster within the *striatum*. We also explored the functions highlighted by the “core LRRK2 interactors” within each tissue and illustrated how the LRRK2 interactomes can be used as a tool to trace the relationship between LRRK2 and specific interactors of interest, here exemplified with a study focused on the LRRK2 interactors belonging to the Rab protein family.

## Introduction

Leucine-rich repeat kinase 2 (LRRK2) is a large (285kDa), multidomain protein. As a member of the ROCO superfamily, LRRK2 contains a Ras-of complex (ROC) GTPase domain, as well as a serine-threonine kinase domain, linked to the ROC domain by a C-terminal-of-ROC (COR) domain of unclear function. This enzymatic core is flanked by 4 protein-protein interaction domains (Guaitoli *et al*, 2016). This complex domain structure makes LRRK2 an interesting enzyme from a biochemical perspective, suggesting it may act as a signalling hub able to orchestrate different cellular functions. Mutations in the *LRRK2* gene are the most common genetic causes of familial Parkinson’s disease (PD), accounting for 2-40% of cases depending on the population under analysis (Drolet *et al*, 2011), with the most common (and most intensively studied) mutation being a pathogenic G2019S amino acid change located in the kinase domain (Healy *et al*, 2008). This mutation has been associated with increased LRRK2 kinase activity, and has been implicated in pathophysiological changes including lysosomal dysfunction, α-synuclein and tau aggregations and dysregulation of neuroinflammation (Tolosa *et al*, 2020). However, although many years of studies have generated a vast array of results, the role(s) of LRRK2 in health and disease still remain elusive.

Apart from PD, multiple lines of evidence have associated LRRK2 with a number of peripheral diseases induced by excessive inflammatory response. *LRRK2* has been identified as a major susceptibility gene for Crohn’s disease (CD) (Michail *et al*, 2013; Umeno *et al*, 2011; Franke *et al*, 2010; Hugot *et al*, 2001; Barrett *et al*, 2008). A newly identified LRRK2-N2081D mutation, which is located in the kinase domain, is associated with increased risk for both PD and CD (Hui *et al*, 2018). Additionally, LRRK2 mutations were reported to aggravate the type-1 reaction in leprosy, and the innate immune response against *mycobacterium tuberculosis* (Fava *et al*, 2016; Weindel *et al*, 2019). These findings indicate a potentially important role of the LRRK2 protein at the interface between the peripheral and the central nervous system (CNS) immunity. Mutations in LRRK2 have also been linked to an increased risk of cancer (Berwick *et al*, 2019). Clinical studies found that PD patients carrying LRRK2-G2019S mutation present higher risk of non-skin cancer, hormone-related cancers and breast cancer compared to non-carriers (Agalliu *et al*, 2015; Saunders-Pullman *et al*, 2010). Although the mechanism underlying the link between LRRK2 and CD, leprosy and cancer is still unclear, this substantial association with peripheral diseases suggests a probably equally important role of the LRRK2 protein in the peripheral tissues as compared to the CNS.

Protein-protein interactions (PPIs) are fundamental for the maintenance of cellular homeostasis, with their alterations (due to mutations or post translational modification) potentially leading to diseases (Lehne & Schlitt, 2009; Vazquez *et al*, 2003). Systems biology deals with the complexity of PPIs applying approaches based on a holistic perspective rather than on a one-to-one perspective. In particular, the core assumption is that proteins interacting with each other constitute a functional unit and thereby are more likely to cooperate in the same cellular pathway(s). In this perspective, the analysis of the protein network built around one or more “seed proteins” of interest allows to gain insights into the biological processes sustained by them, as a community. In this study, LRRK2 was designated as the “seed protein” at the centre of the network analysis, and its interactors were derived from curated peer-reviewed literature. The obtained LRRK2 interactome was then investigated with the goal of describing the cellular functions regulated by LRRK2 together with its interactors under physiological conditions. This strategy had already been employed by 2 comprehensive LRRK2-focused PPI studies which were independently published in 2015 (Manzoni *et al*, 2015; Porras *et al*, 2015). These studies, however, did not investigate the tissue specificity of the LRRK2 interactome.

Multiple lines of evidence have shown that the expression levels of LRRK2 differs greatly in different tissues and cell types, with one possible implication that this variability in expression levels may reflect underlying divergence in the cellular functions of LRRK2 (West *et al*, 2014; Witt *et al*, 2013; Dzamko *et al*, 2013; Kubo *et al*, 2010; Thévenet *et al*, 2011; Taymans *et al*, 2006; Fan *et al*, 2018; Paisán-Ruíz *et al*, 2008). From a cellular perspective, LRRK2 has been associated with many different functions, ranging from modulation of autophagy to control of vesicles dynamics, from regulation of signalling pathways to response to stressors (Jeong *et al*, 2018; Bae & Lee, 2020; Berwick *et al*, 2019; Obergasteiger *et al*, 2019; Manzoni *et al*, 2016, 2013; Cook *et al*, 2017). A way to make sense of this plethora of functions, is to suggest LRRK2 might potentially be implicated in different processes in different tissues, thus reflecting a tissue specific functional profile, which might be a consequence of the existence of tissue specific LRRK2 protein complexes (Lewis & Manzoni, 2012).

Therefore, here we worked on the hypothesis that the interactions between LRRK2 and its partners might be tissue specific. However, since most of PPI data currently available are principally derived from *in-vitro* experiments in cellular models, isolated proteins or from high throughput screening, they lack in tissue specificity. With this work we suggest few different computational approaches to differentiate the general LRRK2 interactome into tissue specific LRRK2 interactomes considering the transcriptomic features and the functional patterns of LRRK2 and its interactors in healthy human tissues. These results provide a tool to model tissue specificity for PPIs and a valuable window onto the role of LRRK2 in health and disease with important implications for the development of safe LRRK2-targeted therapeutic approaches.

## Methods

### Protein-protein interaction (PPI) download

PINOT (http://www.reading.ac.uk/bioinf/PINOT/PINOT_form.html) (Tomkins *et al*, 2020), HIPPIE (http://cbdm-01.zdv.uni-mainz.de/~mschaefer/hippie/index.php) (Alanis-Lobato *et al*, 2017) and MIST (https://fgrtools.hms.harvard.edu/MIST/) (Hu *et al*, 2018) were queried to download “homo sapiens” PPIs for LRRK2 (Uniprot ID: Q5S007, 21 October 2020). To access the broadest possible set of PPI data, “Lenient” filter level was applied in PINOT; while all filters were removed in HIPPIE and MIST.

PPIs were quality controlled via an in-house pipeline described in **Figure 1;** 1) protein IDs of data from different repositories were converted into the same identifier system (HUGO Gene Nomenclature Committee (HGNC) gene names); 2) the “interaction detection methods” in HIPPIE and MIST were reassigned referring to the PINOT Method Grouping Dictionary (Lenient version). The PINOT Method Grouping Dictionary clusters similar detection methods annotated in PSI-MI ontology (e.g. “Two hybrid fragment pooling approach MI:0399” and “Two hybrid bait and prey pooling approach MI:1113” are allocated in the same category: “Two Hybrid”); 3) PPIs extracted from the 3 databases were merged after removing duplicates; 4) LRRK2 interactors were then scored (Final Score, FS) by adding the number of detection methods (Method Score, MS) and the number of reporting publications (Publication Score, PS). The LRRK2 interactome was generated from interactors with FS > 2. Interactors with lower FS (≤ 2) were removed from further analysis because of their poor reliability (either were not replicated in multiple experiments or with missing publication identifier or with missing record of detection method). Of note, interactors marked as “Unreviewed” in UniProtKB were removed as well. PPIs that passed quality control thereby constitute what we defined as the general LRRK2 interactome (LRRK2_int_). Family classification of QC-ed LRRK2 interactors was extracted (on 19 December 2020) from UniProt via the R package: UniProt.ws (Carlson, 2021).

**Figure 1:**
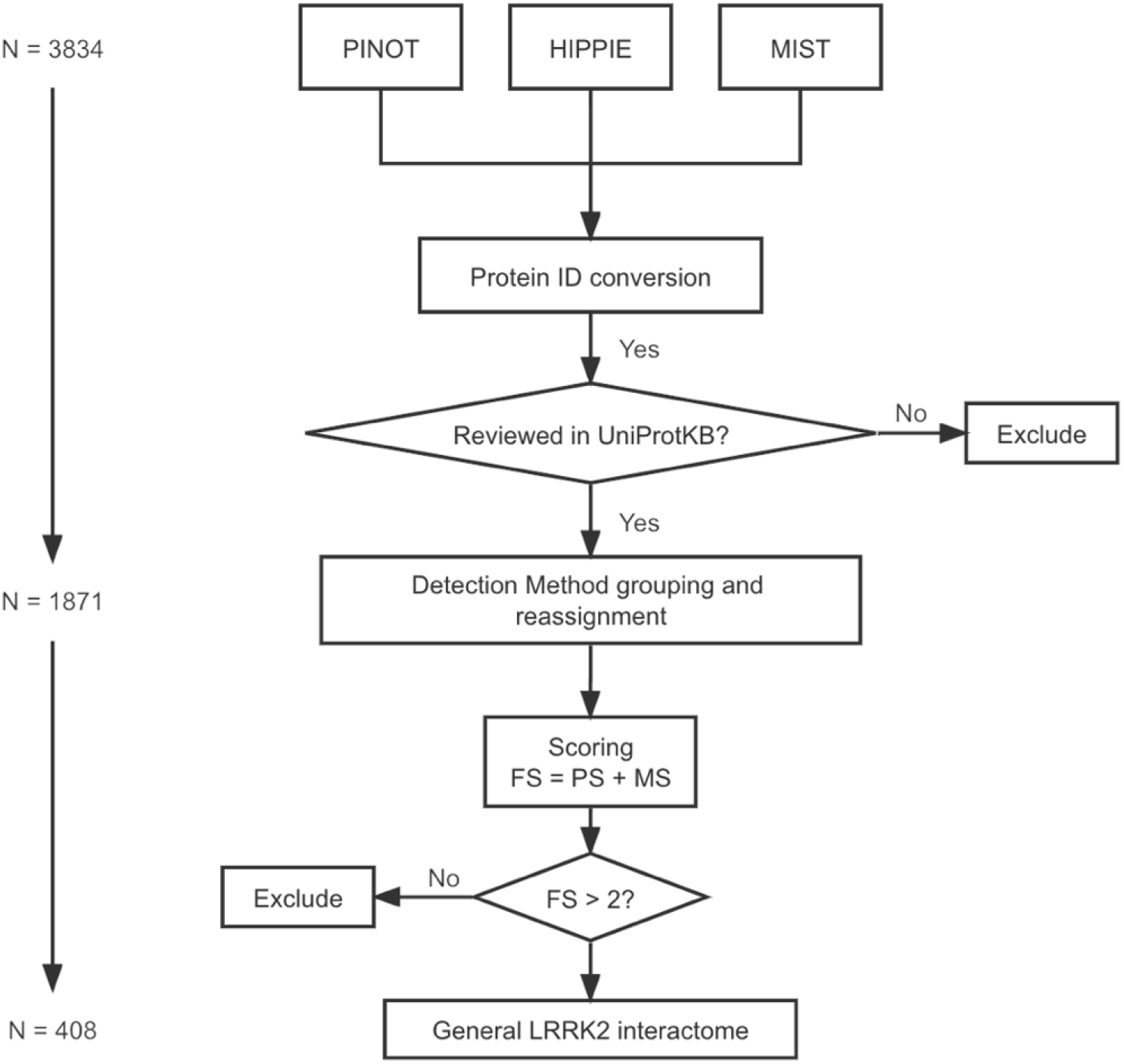
PPI quality control pipeline for LRRK2 interactome construction. Human PPI data downloaded from PINOT, HIPPIE and MIST databases were merged after ID conversion using HGNC gene names. Merged data underwent interaction detection method reassignment using an in-house dictionary. Publication score (PS) was defined as the number of papers in which a PPI was reported, while method score (MS) was defined as the number of different methods by which a PPI was detected. Final score (FS) was calculated as PS + MS. PPIs with FS ≤ 2 were excluded from further analysis.

### Functional characterisation of the general LRRK2_int_

The general LRRK2_int_ was analysed by Functional Enrichment Analysis using the tool g:GOSt on 1 Oct 2021 (g:Profiler (https://biit.cs.ut.ee/gprofiler/gost) (Raudvere *et al*, 2019). The parameters were set as follows: organism - Homo sapiens (Human); data source - GO biological process (GO-BPs) only; statistical domain scope – annotated genes only; statistical method - Fisher’s one-tailed test; significance threshold – Bonferroni correction (threshold = 0.05). No hierarchical filtering was included. To increase the sensitivity of analysis, a cut-off of ≤ 1500 was set for the “term size” of enriched GO terms. Finally, GO-BPs whose enrichment gene set did not include LRRK2 were discarded, thereby keeping only the GO-BPs that LRRK2 directly contributed to. The remaining GO-BPs were grouped according to their semantic similarity (hereby referred as “GO-BP groups”) using an in-house dictionary followed by manual scrutiny. LRRK2 interactors contributing to the enrichment of each GO-BP group were extracted (hereby referred as “functional groups”). The composition (in terms of LRRK2 interactors) of different functional groups was compared using Multiple Correspondence Analysis (MCA).

### RNA-Seq data download and quality control

RNA-seq data (read counts) were downloaded for 56200 genes in 11 brain regions (amygdala, anterior cingulate cortex, caudate (basal ganglia), cerebellum/cerebellar hemisphere, cortex/frontal cortex (BA9), hippocampus, hypothalamus, nuclear accumbens (basal ganglia), putamen (basal ganglia), spinal cord (cervical c-1) and substantia nigra (basal ganglia)), 3 peripheral tissues (lung, liver, kidney) and whole blood from the Genotype-Tissue Expression (GTEx, https://www.gtexportal.org/home/) Analysis Release V8 (dbGaP Accession phs000424.v8.p2) on 19 Aug 2021 (https://storage.googleapis.com/gtex_analysis_v8/rna_seq_data/GTEx_Analysis_2017-06-05_v8_RNASeQCv1.1.9_gene_reads.gct.gz). Of note, in GTEx, “cerebellum/cerebellar hemisphere” and “cortex/frontal cortex (BA9)” are duplicated pairs (https://www.gtexportal.org/home/faq-brainCortexAndCerebellum). Therefore, we kept “cerebellum” and “cortex” in our analysis because they contain more samples compared to their duplicates (209 vs. 175; 205 vs. 175). For kidney, we only kept data for cortex, while data for medulla were discarded because it contains too few samples (n = 4).

RNA-seq data was then quality controlled following an in-house pipeline: for each tissue, 1) low count genes (with read counts = 0 in more than 5% of all samples) were discarded; 2) samples with mapping rates < 80% were discarded (GTEx’s Sample Attribute file: https://storage.googleapis.com/gtex_analysis_v8/annotations/GTEx_Analysis_v8_Annotations_SampleAttributesDS.txt); 3) expression-profile similarity among remaining samples in a given tissue was examined via pairwise Pearson’s correlation test on the read counts; 4) samples were clustered based on the correlation coefficients calculated in 3). Samples that lay outside the main cluster(s) in the hierarchical dendrogram were recognised as “outliers” and were thereby discarded from further analysis. Read counts for LRRK2 interactors were extracted using HUGO gene names and Ensembl Gene IDs. LRRK2 interactors with available expression data for all tissues were kept for further analysis.

### Evaluation of tissue specific LRRK2_ints_ based on differential expression

Pair-wise Differential Expression Analysis (DEA) was performed to compare the expression values of LRRK2 interactors across different tissues via the R package “DESeq2” (Love *et al*, 2014). In “DESeq2”, p-values generated during each round of DEA between every pair of tissues were automatically adjusted for multiple testing correction. These p-values were further corrected using Bonferroni’s method to reduce the Type II error generated from pair-wise DEA among all tissues of analysis. This resulted in a matrix containing DEA p-values for each possible pair of tissues, for all LRRK2 interactors. For each interactor, tissues were ranked according to the DEA results via the following method: for interactor *I*, if its expression level in Tissue *A* is significantly higher than in Tissue B (Bonferroni corrected p-value < 0.05), then 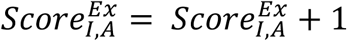, while 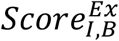 remains unchanged, and vice versa. If the comparison between Tissue *A* and Tissue *B* is insignificant (Bonferroni corrected p-value ≥ 0.05), both 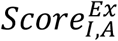 and 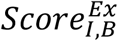 remain unchanged. According to this classification, for interactor 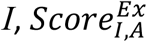 was directly indicative of the expression level of interactor *I* in Tissue *A* in comparison with the other tissues: the higher the 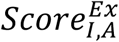, the higher the expression of interactor *I* in Tissue *A*. Interactors that presented uniquely and significantly high expression in certain tissues 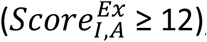, meaning that the expression level of Interactor *I* in Tissue *A* is higher than 85.7% (12/14) tissues.

Finally, a heatmap (Heatmap_DEA) was generated based on the mean expression values of LRRK2 interactors in different tissues calculated from the normalised read counts matrix derived from DEA using R package “gplots” (“DESeq2” transforms the read counts via internal normalisation where geometric mean is calculated for each gene across all samples) (Love *et al*, 2014). Two dendrograms were derived from Heatmap_DEA: 1) hierarchical clustering of tissues based on the similarity of LRRK2_int_’s expression distribution (Den_DEA1); 2) hierarchical clustering of LRRK2 interactors based on the similarity of their expression patterns across different tissues (Den_DEA2). By cutting these 2 dendrograms, we identified 1) clusters of tissues in which the LRRK2_int_ presented similar expression behaviours; 2) clusters of interactors that exhibited similar expression patterns with LRRK2 across different tissues (DEA_Cluster_LRRK2_).

### Evaluation of tissue specific LRRK2-ints based on co-expression analysis

Read counts data derived from GTEx was used to calculate Pearson’s correlation between LRRK2 and its interactors in each single tissue. Multiple testing correction was performed using Bonferroni’s method. The co-expression patterns of LRRK2 with its interactors in each tissue were evaluated by comparison to the corresponding reference co-expression coefficient distribution in the same tissue, which was generated (for each tissue) from co-expression analysis between LRRK2 and 1000 sets of randomly picked genes (to match the size of the general LRRK2_int_) in GTEx.

The distribution of the co-expression coefficients for the LRRK2 interactors was compared across different tissues via the following steps: 1) One-way ANOVA followed by Tukey’s test was performed to compare the coefficients across tissues; 2) if the co-expression coefficients are significantly higher in Tissue *A* than in Tissue *B* (adjusted p-value < 0.05), then 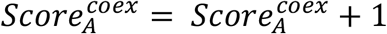, while 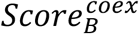 remains unchanged. If the comparison between Tissue *A* and Tissue *B* is insignificant (adjusted p-value ≥ 0.05), both 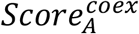 and 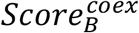 remain unchanged. According to this method, for interactor *I*, the 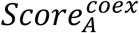 in was directly indicative of the co-expression level of interactor *I* with LRRK2 in Tissue *A*.

Finally, a heatmap (Heatmap_Co-ex) was generated based on the matrix of co-expression coefficients between interactors and LRRK2 in different tissues using the R package “gplots”. Two dendrograms were derived from Heatmap_Co-ex: 1) hierarchical clustering of tissues based on the similarity of LRRK2-interactor coefficient distribution (Den_Co-ex1); 2) hierarchical clustering of LRRK2 interactors based on the similarity of their co-expression (with LRRK2) behaviours across different tissues (Den_Co-ex2). By cutting the 2 dendrograms, we identified 1) clusters of tissues in which the LRRK2_int_ presented similar co-expression behaviours; 2) clusters of interactors that exhibited similar co-expression patterns with LRRK2 across different tissues. In addition, we compared the co-expression coefficients of the interactor clusters identified in 2) via t-test, thereby identifying the cluster of interactors that presented the highest co-expression with LRRK2 (Co-ex_Cluster_LRRK2_).

### Tissue-specific LRRK2 interactomes and functional patterns

Tissue specific LRRK2_ints_ were constructed by combining the results of DEA and co-expression analysis. In each tissue specific LRRK2_int_, core interactors were defined as LRRK2 interactors that i) either presented significantly higher expression in the given tissue as compared to other tissues of analysis, or ii) exhibited high co-expression in the given tissues in comparison to other interactors in the LRRK2_int_. The “highly expressed” interactors were filtered via the following steps: 1) the absolute expression levels (automatically transformed read counts generated by the R package “Deseq2”) were log transformed to reach a normal distribution; 2) threshold of “high expression” was set as 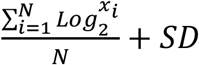 (*N*: the total number of LRRK2 interactor in the tissue specific LRRK2_int_, *SD*: standard deviation of transformed expression levels of LRRK2 interactors in the given tissue); 3) interactors with transformed expression values above the threshold were defined as “highly expressed” interactors in the given tissue. Highly co-expressed interactors were defined via the following steps: for each tissue, 1) LRRK2:interactor co-expression coefficients were transformed via the formula 10^*μ*^ (*μ* = co-expression coefficient) to reach a normal distribution; 2) threshold of “high co-expression” was set as 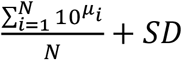 (*N*: the total number of LRRK2 interactor in the tissue specific LRRK2_int_, *SD*: standard deviation of transformed co-expression coefficients between LRRK2 and its interactor in the given tissue); 3) interactors with transformed co-expression coefficients above the threshold were defined as “highly co-expressed” interactors in the given tissue. We analysed functional pattern of each tissue specific LRRK2_int_ by analysing the biological processes in which its core interactors were involved, based on the assumption that interactors that are higher expressed or co-expressed with LRRK2 are more likely to contribute to cellular functions together with LRRK2.

## Results

### Construction of the general LRRK2_int_

A total of 1436, 548 and 1850 human LRRK2 interactors were retrieved from PINOT, HIPPIE and MIST respectively (**Figure 1**). To harmonise different protein identifiers, all protein IDs were converted to HUGO gene names. One interactor (Entrez ID: 333931) was removed because its record is no longer valid in NCBI Gene. Furthermore, 3 proteins coded by transcriptional read-throughs (RPL17-C18orf32, TPTEP2-CSNK1E and BUB1B-PAK6) were marked as “Unreviewed” entries in UniProtKB, and were thereby discarded from further analysis. After protein ID conversion, the 3 protein sets (3831 annotations in total) were merged into 1 list of 1871 unique interactors (hereby referred as “merged list”), suggesting that albeit the differences in data sources and versions, there is a large amount of overlap among PINOT, HIPPIE and MIST in terms of PPIs retrieved for LRRK2. Of note, among the 1871 interactors, 529 (28.3%) were shared by all the 3 tools; 902 (48.2%) were downloaded from 2 of the 3 databases; 440 (23.5%) were present in only 1 database (10 in PINOT, 11 in HIPPIE and 419 in MIST).

For each PPI in the merged list, the “interaction detection methods” were extracted and reassigned according to PINOT Method Grouping Dictionary (Lenient version), where similar methods are grouped together. In this way only technically different interaction detection methods were considered as independent evidence of protein interaction, thus adding stringency to the PPI collection pipeline.

The LRRK2 interactors were scored based on the number of publications (Publication Score, PS) and different types of detection methods (Method Score, MS), thereby generating the Final Score (FS = PS + MS). A total of 1463 “low-quality” interactors (1463/1871, 78.2%) with an FS ≤ 2 (indicating that these LRRK2 interactions were reported in 1 publication and with 1 method only, therefore never replicated) were identified and removed from further analyses. A final list containing 408 LRRK2 interactors with FS > 2 was obtained (hereby referred as “the general LRRK2_int_”, **Table S1**). Among the 408 interactors, 352 (86%) were scored FS ≤ 5; 41 (10%) were scored between 6 and 8; 18 (4%) were scored FS ≥ 9. LRRK2 itself exhibited the highest FS = 50, as many publications confirmed LRRK2 as able to self-interact. Other robust LRRK2 interactors were HSP90AA1 (FS = 19); YWHAQ/14-3-3T (FS = 14), followed by HSPA8, MSN, YWHAZ/14-3-3Z, CDC37, DNM1L, STUB1 and TUBB (**Figure 2A**).

**Figure 2:**
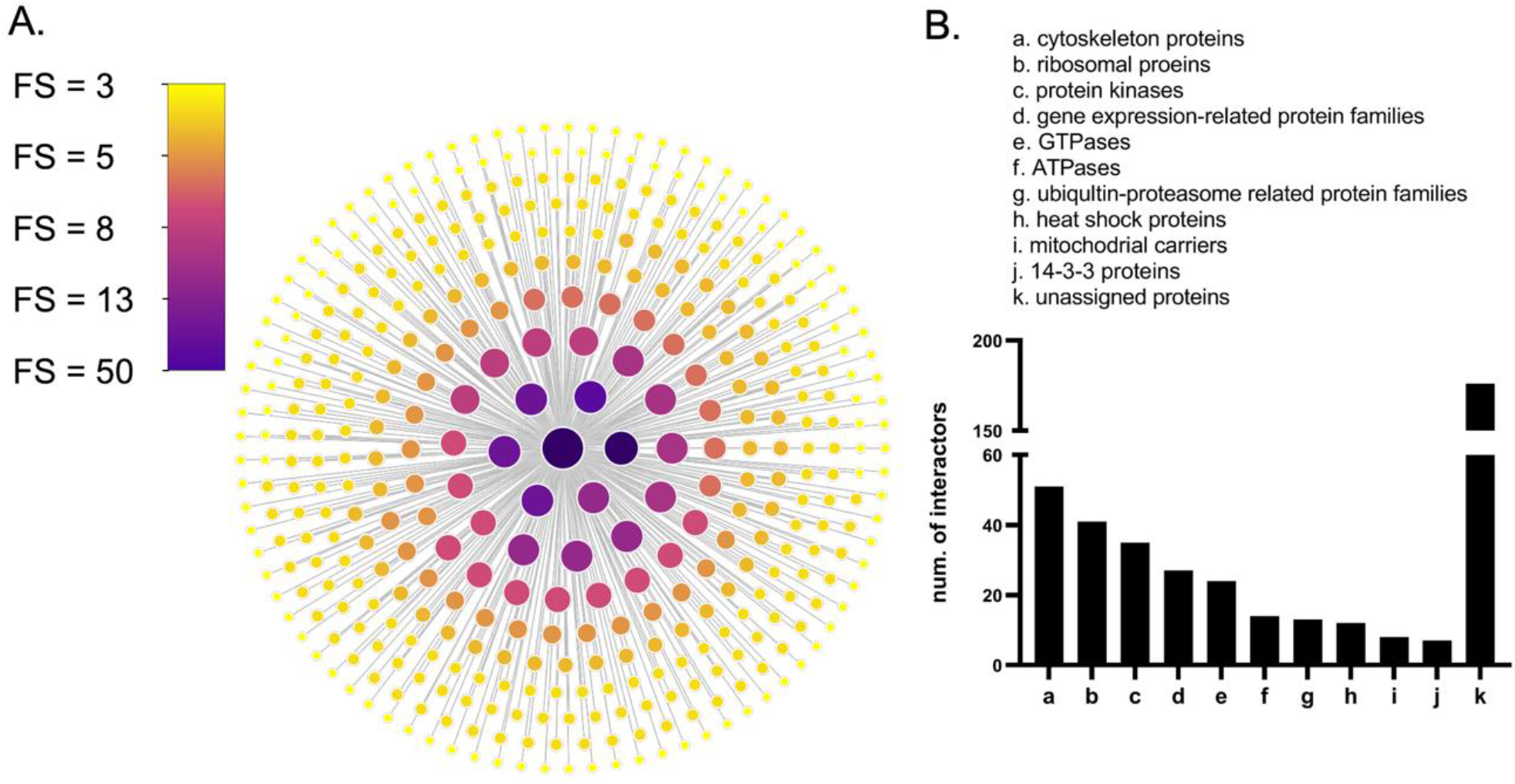
The general LRRK2 interactome. A) Nodes (n = 408) represent LRRK2 interactors that were with FS > 2 and were reviewed by the UniprotKB. Node fill colour and node size are weighted on the final score (FS). Larger size and darker colour indicate higher FS. B) Family classification of LRRK2 interactors.

Out of the 408 LRRK2 interactors, 232 (56.9%) were classified into families based on UniProt record: cytoskeleton proteins (n = 51 interactors, 12.5%), ribosomal proteins (n = 41 interactors, 10.0%), protein kinases (n = 35 interactors, 8.6%), GTPases (n = 24 interactors, 6.6%), ATPases (n = 14 interactors, 3.4%), heat shock proteins (n = 12 interactors, 2.9%), and mitochondrial carriers (n = 8, 2.0%). In addition, 13 (3.2%) LRRK2 interactors were classified in ubiquitin-proteasome related protein families; while 27 interactors (6.7%) belonged to gene expression-related families (8 transcription factors/regulators, 5 helicases, 4 splicing factors and 10 DNA-metabolism-related proteins) (**Figure 2B**). Of note, the LRRK2 interactors included 13 Rab GTPases and seven 14-3-3 proteins. The Rab GTPase family and the 14-3-3 family are the 2 groups of proteins that have been most widely recognised as LRRK2 interactors (Mason *et al*, 2009; Seol *et al*, 2019).

### Functional enrichment analysis on the general LRRK2_int_

Functional Enrichment Analysis was performed for the general LRRK2_int_. A total of 597 significant GO-BPs (Bonferroni adjusted p-values < 0.05) were returned. A cut-off of “term size” ≤ 1500 was applied to the enriched GO-BPs to remove general terms (140/597, 23.4% of terms removed) and improve sensitivity of analysis. GO-BPs whose enrichment gene set (i.e. intersection) did not contain LRRK2 were also removed leading to the final enrichment list containing 183 GO-BPs. These GO-BPs were clustered into the following 14 functional groups based on their semantic similarities: “autophagy”, “cell death”, “development, “intracellular organisation”, “immune system”, “metabolism”, “protein catabolism”, “protein localisation”, “protein modification”, “response to stress”, “regulation of enzyme functions”, “regulation of gene expression”, “signalling”, and “transport”.

LRRK2 interactors contributing to the enrichment of each single functional group were extracted. A total of 315 (77.2%) LRRK2 interactors contributed to the enrichment of at least 1 functional group, while 10 LRRK2 interactors contributed to the enrichment of ≥ 13 out of the 14 functional groups (AKT1, CDK5, GSK3B, PRKCZ, HSP90AB1, HSP90AA1, MAPT, HIF1A, HDAC6, TP53), suggesting their interactions with LRRK2 are involved in multiple biological processes. The largest numbers of LRRK2 interactors were found to contribute to the functional groups of: “intracellular organisation” (n = 192/315 interactors, 60.7%), “transport” (n = 161/315 interactors, 50.9%) “metabolism” (n = 160/315 interactors, 50.6%), “regulation of gene expression” (n = 147/315 interactors, 46.5%) and protein catabolism (n = 137/315 interactors, 43.4%), confirming the observation already made for the most represented functional groups (**Figure 3A**).

**Figure 3:**
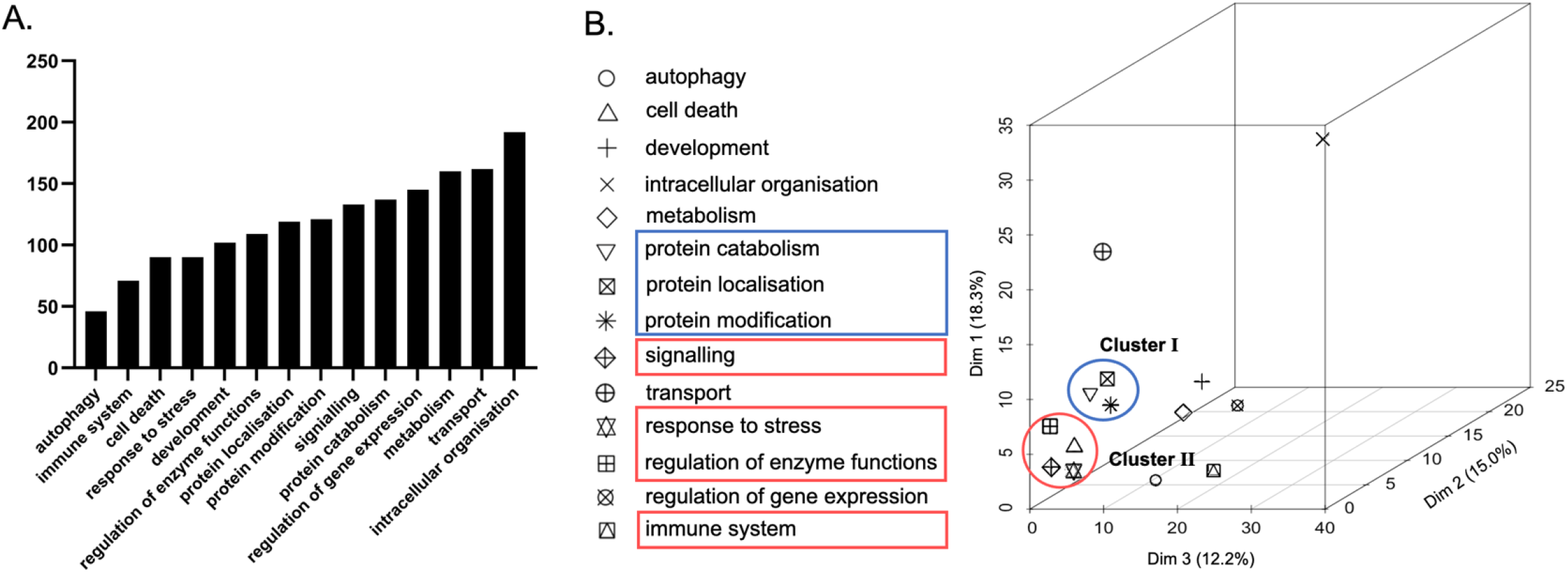
Functions of LRRK2_int._ A) The bar graph shows the numbers of interactors contributing to the enrichment of different functional groups. B) The 14 functional groups were then analysed via MCA to compare their composition in terms of LRRK2 interactors. Two clusters of functional groups were identified, suggesting each of these clusters comprises similar LRRK2 interactors.

Multiple Correspondence Analysis (MCA) was performed to compare the composition (in terms of LRRK2 interactors) of the 14 functional groups (**Figure 3B**). The top three components (Dim.1, Dim.2, Dim.3) accounted for 35.8% of total variability in the data (18.6%, 15.0% and 12.2%, respectively). “Development”, “intracellular organisation”, “metabolism”, “protein localisation” and “transport” presented as independent elements as they were distant from every other functional group in the MCA graph. “Protein localisation”, “protein modification” and “protein catabolism” were clustered in Cluster I, while “signalling”, “regulation of enzyme functions”, “immune system” and “response to stress” were clustered in Cluster II. This suggested the functional groups within each of the two clusters have a similar composition in terms of LRRK2 interactors.

### RNA-Seq data download and quality control

RNA-Seq read counts data of 56200 genes for 11 brain regions, lung, liver, kidney (cortex) and whole blood were downloaded from GTEx. More than 50% of the genes were discarded due to low read counts (identified as having more than 5% of samples with null read counts in a given tissue). Of note, a total of 7 LRRK2 interactors were removed in this step, reducing the dimension of the LRRK2_int_ from 408 to 401 analysable interactors. Pairwise Pearson’s correlation test was performed within each tissue to evaluate the similarity of gene expression profiles read count across the samples. Samples were grouped by hierarchical clustering based on the Pearson’s coefficients to identify samples whose gene expression profile was generally off-scale. No outlier samples were found in any tissue, suggesting that GTEx RNA-Seq data are barely affected by batch effects. Read counts for LRRK2 interactors were then extracted using HUGO gene names and Ensembl Gene IDs. Five proteins were not found in GTEx Ensembl Gene ID list and were thereby removed. Seventeen LRRK2 interactors presented missing expression values in certain tissues and were also excluded. In total, read counts for 379 (92.9%) LRRK2 interactors were obtained for further analysis.

### Construction of tissue specific LRRK2_ints_

#### Differential Expression analysis on the LRRK2_int_

Pair-wise differential expression analyses (DEA) were performed for each LRRK2 interactor across different tissues. Tissues were ranked based on the DEA results of each LRRK2 interactors. We were able to observe the following tissue-specific expression patterns of the general LRRK2_int_: i) the tissues with the largest numbers of LRRK2 interactors showing significantly high expression levels are blood (with 163 LRRK2 interactors expressed significantly higher in blood than in ≥ 10 of the other tissues analysed 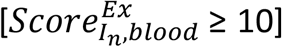), cerebellum, spinal cord c-1 and frontal cortex (with n = 117; n = 74; and n = 67 highly expressed LRRK2 interactors respectively, all with 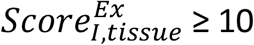). The expression levels of LRRK2 interactors were generally lower in caudate, putamen, kidney cortex and liver (scored < 9 in 98.1% of all interactors) (**Figure 4A**). When considering the individual components of the LRRK2_int_, a total of 185 out of 379 interactors (48.8%) presented with unique and significantly high expression in a certain tissue (with a 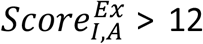, suggesting that the expression level of Interactor *I* is significantly higher in Tissue *A* than in ≥ 12 other tissues analysed). These 185 highly expressed interactors were distributed across 6 tissues (cerebellum, frontal cortex, spinal cord c-1, hypothalamus, anterior cingulate cortex and blood) and showed high tissue-specificity, i.e. they were highly expressed only in 1 tissue (**Figure 4B**).

**Figure 4.**
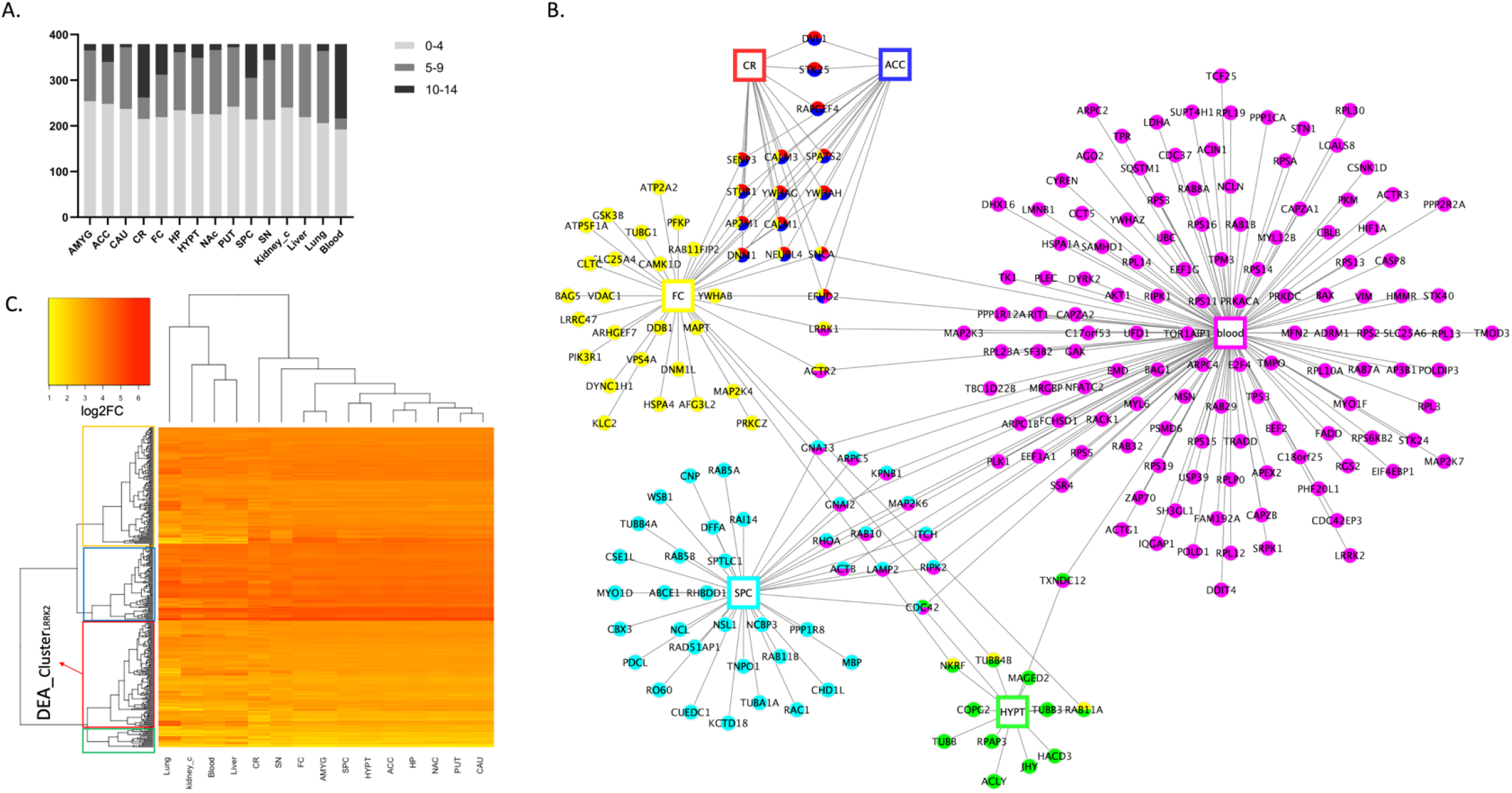
Differential Expression Analysis (DEA) on the LRRK2_int_. A) DEA was performed to compare the expression levels of each LRRK2 interactor across different tissues. Tissues were ranked based on the significant comparison results. The bar graph shows the distribution of the ranks of different tissues. Each bar in the graph represents a tissue, and segments in the bar represent ranks of that tissue at three levels: AVERAGE (≤ 4); MIDDLE (between 5 and 9); HIGH (≥ 10). Of note, an AVERAGE rank suggests that for interactor I, the expression level in a given tissue is lower/not significantly higher than in other tissues, or only higher than in ≤ 4 tissues (< 27% of all tissues), while a HIGH rank means the expression level of interactor I is significantly higher in a given tissue than in ≥ 10 other tissues (> 67% of all tissues). B) The network graph shows the LRRK2 interactors with significantly high expression in certain tissues (tissue ranks ≥ 12), suggesting that the expression levels of these interactors in a specific tissue are significantly higher than in ≥ 12 other tissues (86% of all tissues). Tissues are represented as rectangular nodes, while interactors are represented as round nodes. Different colour indicates different tissues. C) Heatmap_DEA was generated from normalised read counts (log2 transformed) of LRRK2 interactors in different tissues derived from DEA. Darker colour represents higher expression levels. The horizonal dendrogram of Heatmap_DEA was extracted as Den_DEA1. It shows the hierarchical clustering of tissues in which the LRRK2_int_ exhibits similar expression patterns. The vertical dendrogram of Heatmap_DEA was extracted as Den_DEA2. It shows the hierarchical clustering of LRRK2 interactors based on the similarity of their expression figures across different tissues. Den_DEA2 was cut to generate 4 clusters of LRRK2 interactors (Cluster 1-4, marked in green, red, blue and yellow, respectively). The cluster containing LRRK2 (marked in red) is defined as DEA_ClusterLRRK2, in which the interactors presented similar overall expression distribution across tissues as LRRK2. Abbreviations: ACC: Anterior Cingulate Cortex; AMYG: Amygdala; CAU: caudate; CR: cerebellum; FC: frontal cortex; HP: hippocampus; HYPT: hypothalamus; NAc: nucleus accumbens; PUT: putamen; SN: substantia nigra; SPC: spinal cord c-1; Kidney_c: kidney cortex.

A heatmap was generated based on the normalised expression matrix derived from DEA (Heatmap_DEA, **Figure 4C**). Two dendrograms were extracted from Heatmap_DEA: 1) Den_DEA1 for the hierarchical clustering of tissues based on the overall LRRK2_int_ expression patterns; 2) Den_DEA2 for the hierarchical clustering of each LRRK2 interactor based on expression behaviours across different tissues. In Den_DEA1, brain regions and peripheral tissues presented in two distinct groups, suggesting that the overall expression levels for the components of the LRRK2_int_ are different in the brain in comparison with other tissues. Among the 11 brain regions, putamen, caudate, and nucleus accumbens were clustered together, indicating that the LRRK2 interactors exhibited similar expression patterns in these 3 brain regions (in terms of absolute expression levels). Of note, putamen, caudate and nucleus accumbens regions are the fundamental parts of striatum, which is substantially impacted with PD pathology. Den_DEA2 was cut following the principle of obtaining the largest number of clusters while avoiding generating clusters comprising single interactors, thereby 4 clusters were obtained (Cluster 1-4; n = 25, 124, 91 and 138 interactors, respectively). A total of 124 interactors presented in the same cluster as LRRK2 (Cluster 1, hereby referred as DEA_Cluster_LRRK2_), suggesting these proteins exhibited a similar expression pattern (in terms of absolute expression levels) when compared with LRRK2 in different tissues.

#### Co-expression analysis on the general LRRK2_int_

Pearson’s correlation test was performed to calculate the co-expression coefficients between LRRK2 and each of its 379 interactors in different tissues. The distribution of the 379 co-expression coefficients for each tissue was compared to the corresponding reference distribution generated from co-expression analysis between LRRK2 and 1000 sets of randomly picked genes (for each random gene list, n = 379 to match with the dimension of the general LRRK2_int_). In frontal cortex, putamen, nucleus accumbens, hypothalamus, anterior cingulate cortex, caudate, and cerebellum, LRRK2_int_ presented a larger distribution of high co-expression coefficients (0.65 to 0.85) in comparison to the reference, indicating that LRRK2 interactors were strongly correlated with LRRK2 in comparison with randomly picked genes. In hippocampus, spinal cord c-1 and lung, the LRRK2 interactors presented more moderate co-expression with LRRK2 (0.45 to 0.65) in comparison to the reference, suggesting they are mildly correlated with LRRK2 in comparison with randomly picked genes. The distribution of co-expression coefficients of the LRRK2_int_ overlapped with the reference for amygdala, substantia nigra, blood, liver and kidney cortex, suggesting no significant co-expression between LRRK2 and its interactors was present in these tissues.

One-way ANOVA followed by Tukey’s test was performed to compare the distribution of co-expression coefficients across tissues. The tissues were then ranked based on the significant results. Tissues with the higher co-expression coefficients between LRRK2 and its interactors were putamen (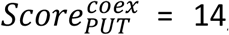, suggesting that the co-expression coefficients were significantly higher in putamen compared to all other 14 tissues analysed) and nucleus accumbens 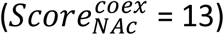, followed by caudate and hypothalamus 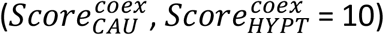. Hippocampus was the brain region with the lowest number of interactors co-expressed with LRRK2 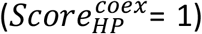. Kidney cortex obtained the highest rank among the peripheral tissues 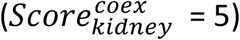 (**Figure 5A**).

**Figure 5:**
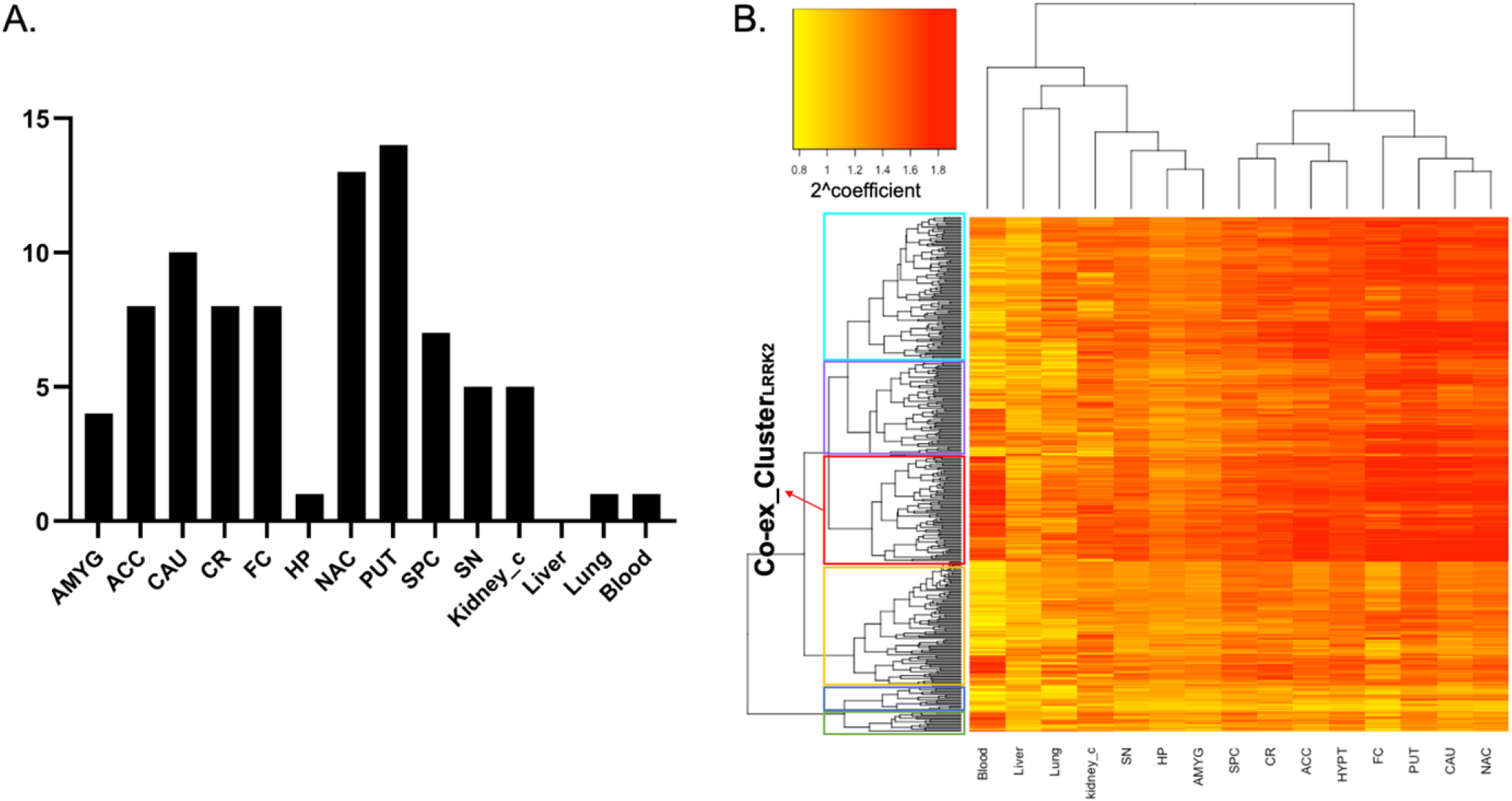
Co-expression analysis on the LRRK2_int_. A) Pair-wise Tukey’s test was performed to compare the co-expression coefficients (interactors vs LRRK2) across different tissues. Tissues were ranked according to the results. The bar graph shows that putamen, nucleus accumbens, caudate and hypothalamus are tissues with the highest ranks. Liver presents a rank of 0, meaning the co-expression coefficients of LRRK2 interactors are the lowest in comparison with any other tissues analysed. B) The heatmap was generated from the coefficient matrix derived from the co-expression analysis (Heatmap_Co-ex). Darker colour represents higher co-expression coefficient. The horizonal dendrogram of Heatmap_Co-ex was extracted as Den_Co-ex1, which shows the hierarchical clustering of tissues in which the LRRK2 interactors exhibited similar co-expression patterns with LRRK2. The vertical dendrogram of Heatmap_Co-ex was extracted as Den_Co-ex2, which shows the hierarchical clustering of interactors based on the similarity of their co-expression figures with LRRK2 across different tissues. Den_Co-ex2 was cut to generate 6 clusters of LRRK2 interactors (Cluster A-F, marked in green, blue, yellow, red, purple and turquoise, respectively). Interactors in Cluster D presents the highest level of overall co-expression behaviour with LRRK2 across different tissues (referred as Co-ex_Cluster_LRRK2_). Abbreviations: ACC: Anterior Cingulate Cortex; AMYG: Amygdala; CAU: caudate; CR: cerebellum; FC: frontal cortex; HP: hippocampus; HYPT: hypothalamus; NAc: nucleus accumbens; PUT: putamen; SN: substantia nigra; SPC: spinal cord c-1; Kidney_c: kidney cortex.

A heatmap was generated from the coefficient matrix derived from co-expression analysis (Heatmap_Co-ex, **Figure 5B**). The 2 dendrograms in Heatmap_DEA were extracted as follows: 1) Den_Co-ex1 for the hierarchical clustering of tissues in which the LRRK2_int_ presented similar co-expression patterns; 2) Den_Co-ex2 for the hierarchical clustering of the LRRK2 interactors that exhibited similar co-expression behaviours with LRRK2 across different tissues. In Den_Co-ex1, two clusters were identified: i) frontal cortex, putamen, nucleus accumbens and caudate; ii) spinal cord c-1, hypothalamus, cerebellum and anterior cingulate cortex, indicating that the pattern of co-expression between LRRK2 and its interactors was different between the 2 clusters but similar for the tissues within each cluster. This suggests that the LRRK2_int_ may participate in different cellular functions in the 2 clusters of brain regions.

A cut-off was applied to the Den_Co-ex2 to cluster LRRK2 interactors based on the similarity of co-expression behaviour (with LRRK2) across different tissues. The cut-off was set to obtain the largest number of clusters while avoiding generating clusters comprising single interactors. A total of 6 clusters was generated (Cluster A-F; n = 15, 19, 91, 77, 72, 104, respectively), in which Cluster D (n = 77/379, 20.3%) contained the LRRK2 interactors presenting the highest overall co-expression coefficients with LRRK2 across different tissues. Cluster D represents the group of LRRK2 interactors whose genes are more frequently co-expressed with LRRK2 considering the pool of tissues under investigation (hereby referred as Co-ex_Cluster_LRRK2_).

#### Tissue specific LRRK2 interactomes

The DEA and co-expression results were combined to generate 15 tissue specific LRRK2_ints_, which presented distinct expression/co-expression profiles across different tissues. In putamen, caudate and nucleus accumbens, LRRK2 interactors presented moderate expression levels (in comparison with the other tissues analysed) but a larger number of interactors showing high co-expression levels with LRRK2. In frontal cortex, substantia nigra and cerebellum, some interactors exhibited especially high expression levels (in comparison with the other tissues analysed), but the LRRK2:interactors co-expression was generally lower as compared to the 3 brain regions discussed above. Blood was the tissue with the largest number of LRRK2 interactors with high expression levels, but these interactors presented a generally low co-expression with LRRK2. Finally, the LRRK2 interactors presented the lowest expression levels and co-expression (with LRRK2) in hippocampus, liver, lung, and kidney (**Figure 6**).

**Figure 6:**
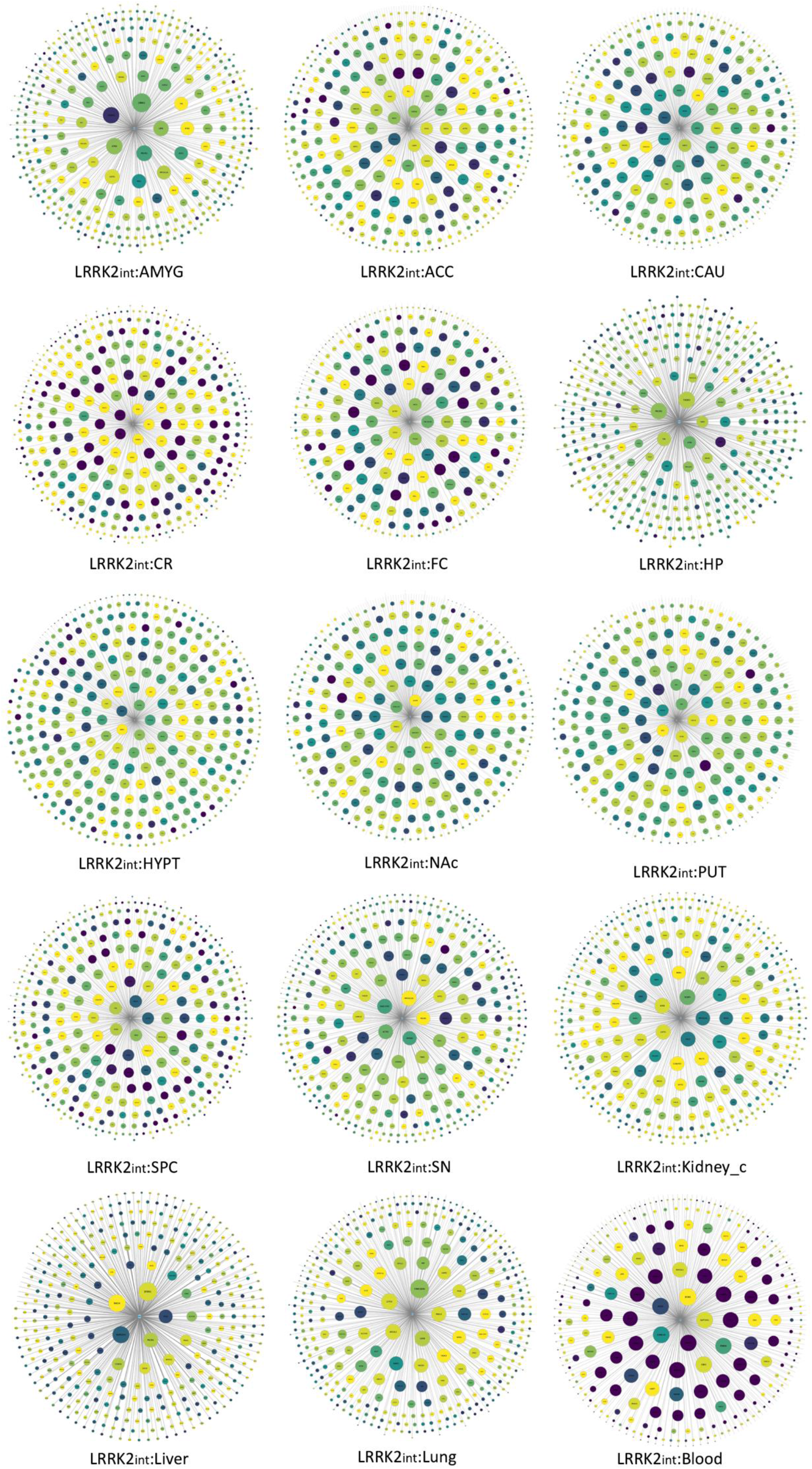
Tissue specific LRRK2_ints._ The network graphs represent the 15 tissue specific LRRK2_ints_ generated from the results of DEA and co-expression analysis. Node colour represents the normalised expression level of LRRK2 interactors. The darker the colour, the higher the expression level of the interactor in a given tissue. Node size represents co-expression coefficient. The larger the node, the higher the co-expression coefficient calculated between LRRK2 and the interactor in a given tissue. Abbreviations: ACC: Anterior Cingulate Cortex; AMYG: Amygdala; CAU: caudate; CR: cerebellum; FC: frontal cortex; HP: hippocampus; HYPT: hypothalamus; NAc: nucleus accumbens; PUT: putamen; SN: substantia nigra; SPC: spinal cord c-1; Kidney_c: kidney cortex.

#### Tissue specific functional patterns of LRRK2_ints_

In order to identify potential functional patterns of the LRRK2_int_ in different tissues, for each tissue, we extracted “core interactors” that either 1) presented high expression levels or 2) exhibited high co-expression behaviours with LRRK2. The functional pattern of LRRK2_int_ of each tissue was obtained by calculating the contribution of the core interactors (for that tissue) to the enrichment of the 14 functional groups identified in the functional enrichment analysis for the general LRRK2_int_ (**Figure 7**).

**Figure 7:**
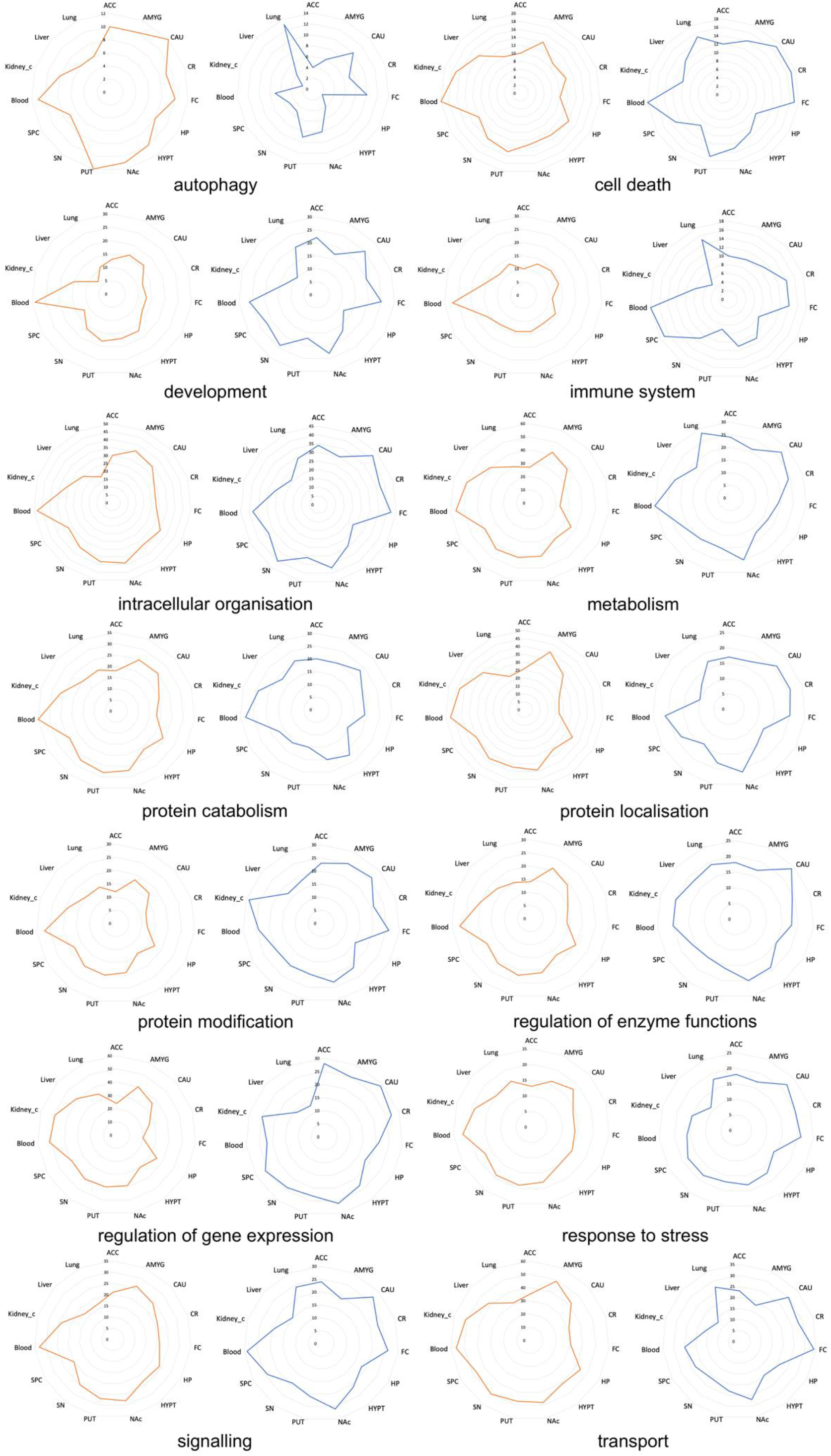
Functional patterns of tissue specific LRRK2_ints._ The radar graphs represent the functional patterns of 15 tissue specific LRRK2_ints_ based on their core interactors. Functional patterns described by highly expressed LRRK2 interactors were represented via orange curve; while the patterns described by highly co-expressed interactors (with LRRK2) were represented via blue curve. Numbers in the graphs represent the numbers of core interactors for different tissues that were included in a given functional group identified in the functional enrichment analysis for the general LRRK2_int_. Abbreviations: ACC: Anterior Cingulate Cortex; AMYG: Amygdala; CAU: caudate; CR: cerebellum; FC: frontal cortex; HP: hippocampus; HYPT: hypothalamus; NAc: nucleus accumbens; PUT: putamen; SN: substantia nigra; SPC: spinal cord c-1; Kidney_c: kidney cortex.

In terms of the brain regions, core interactors in caudate presented high involvement in all 14 functional groups from the perspectives of both highly expressed and co-expressed (with LRRK2) interactors. Core interactors in frontal cortex and nucleus accumbens showed a similar high-engagement pattern in most of the functional groups but only from the perspective of interactors with high co-expression behaviour with LRRK2. For cerebellum, core interactors that presented high co-expression levels with LRRK2 were mainly found contributing to the enrichment of “cell death”, “protein localisation” and “regulation of gene expression”; while in substantia nigra, the core interactors were involved in the functions of “intracellular organisation”, “protein catabolism” and “transport”. Of note, putamen had its highly expressed core interactors engaged in “autophagy”; while core proteins in spinal cord c-1 were primarily participating in the enrichment of the functional group “immune system”; while the core interactors for amygdala were mostly included in the groups of “protein modification”.

As for the peripheral tissues, the core interactors for blood presented involvement in all 14 functional groups. Lung had most of its core interactors involved in “autophagy”, “immune system”, “development” and “metabolism”; while the core interactors in kidney cortex were mainly involved in “metabolism”, “protein modification”, “protein localisation”, “regulation of gene expression” and “transport” mostly from the perspective of highly co-expressed LRRK2 interactors. However, no functional groups were found to include core interactors from liver.

### Application of tissue specific LRRK2_ints_: Differentiating LRRK2:Rab interactions in the CNS and the periphery

The LRRK2_ints_ generated in this work can be used to explore specific interactors of interest; here we selected the Rab proteins as an example to illustrate how the tissue specific LRRK2_ints_ can be filtered to investigate the features LRKR2:Rab protein interaction across different tissues. A total of 13 Rab proteins were identified in the LRRK2_int_ (RAB38, RAB10, RAB11A, RAB11B, RAB11FIP2, RAB1A, RAB1B, RAB29, RAB32, RAB5A, RAB5B, RAB7A and RAB8A), while RAB38 was excluded due to its incomplete expression data across 15 tissues of analysis, thereby only 12 Rab proteins were included for further analysis. Of note, among these Rab proteins, RAB29, RAB8A and RAB11FIP2 presented in the DEA_Cluster_LRRK2_, suggesting these 3 proteins have a similar expression pattern as LRRK2 across all the 15 tissues analysed; while RAB10, RAB11A, RAB11FIP2, RAB1A, RAB1B, RAB5A, RAB5B, RAB7A and RAB8A presented in the Co-ex_Cluster_LRRK2_, indicating that these Rab proteins show a generally higher co-expression with LRRK2 in all the 15 tissues analysed.

All of the 12 Rab proteins contributed to the enrichment of the functional group of “transport” together with LRRK2, while 11 of them exhibited in the group of “intracellular organisation” together with LRRK2, suggesting that these are the 2 functions that are generally sustained by LRRK2-Rab cooperation (**Figure 8**). Five Rab proteins (RAB1A, RAB1B, RAB5A, RAB7A and RAB8A) presented in the “autophagy” functional group, potentially indicating these are the Rab proteins to participate in the waste-disposal processes regulated via LRRK2. Only a small number of Rab proteins was included in the functions of “cell death, protein catabolism, protein modification and regulation of enzyme (≤ 2), though these were the functions presenting the highest enrichment of the general LRRK2 interactors.

**Figure 8:**
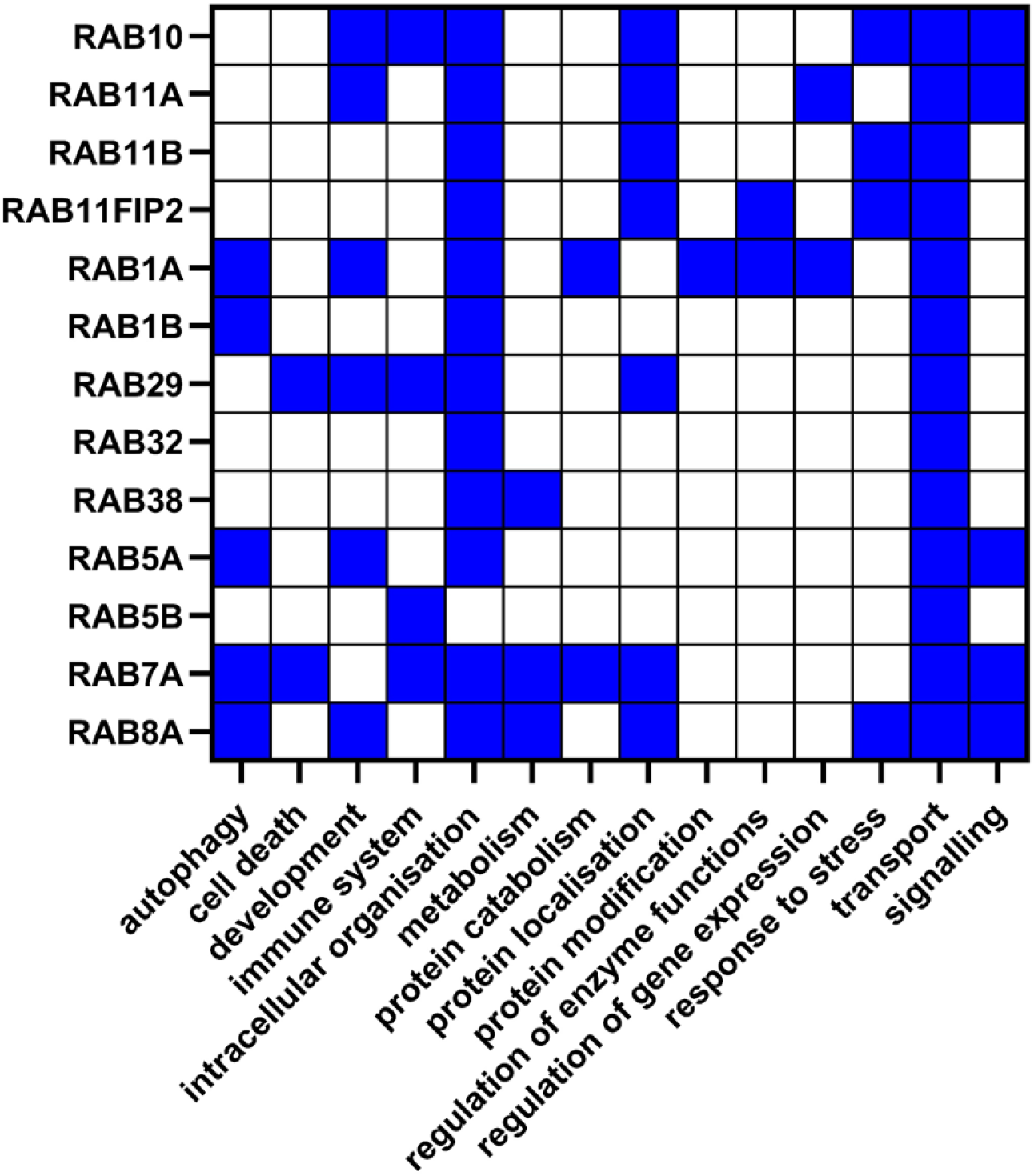
Functional roles of Rab interactors of LRRK2. The heatmap shows the functional groups that included the Rab proteins presented in the LRRK2_int_. Blue squares represent the presence of a certain Rab interactor in a given functional group identified in the functional enrichment analysis for the general LRRK2_int_.

After filtering out non-Rab proteins from the LRRK2_ints_ for each tissue, 15 tissue specific LRRK2:Rab interactomes were generated (**Figure 9**). Overall, RAB7A and RAB5B presented the highest expression level across the majority of tissues analysed, while RAB10, RAB11FIP2, RAB11A, RAB7A and RAB5B exhibited the highest co-expression with LRRK2 in most of the tissues except for amygdala, hippocampus, kidney cortex, lung and liver. Among the brain regions, putamen, caudate, frontal cortex and nucleus accumbens showed identical distribution of absolute expression levels of Rab interactors and similarly high co-expression between these proteins and LRRK2. Of note, RAB32 and RAB29 presented a similar co-expression pattern with LRRK2 across different tissues as compared to other Rab proteins. A high LRRK2:RAB32 was seen in hypothalamus, cerebellum, substantial nigra spinal cord c-1 and blood. Of note, RAB32 presented the highest expression level in blood 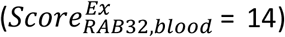, suggesting a potentially more important role of LRRK2:RAB32 interaction in blood. Similarly, RAB29 presented a high co-expression with LRRK2 in hypothalamus, substantia nigra and spinal cord c-1. These suggest a potential co-function among LRRK2, RAB32 and RAB29 in these brain regions. In comparison, LRRK2:Rab co-expression was weak in amygdala and lung, while the poorest co-expression was seen in hippocampus, liver and kidney cortex.

**Figure 9:**
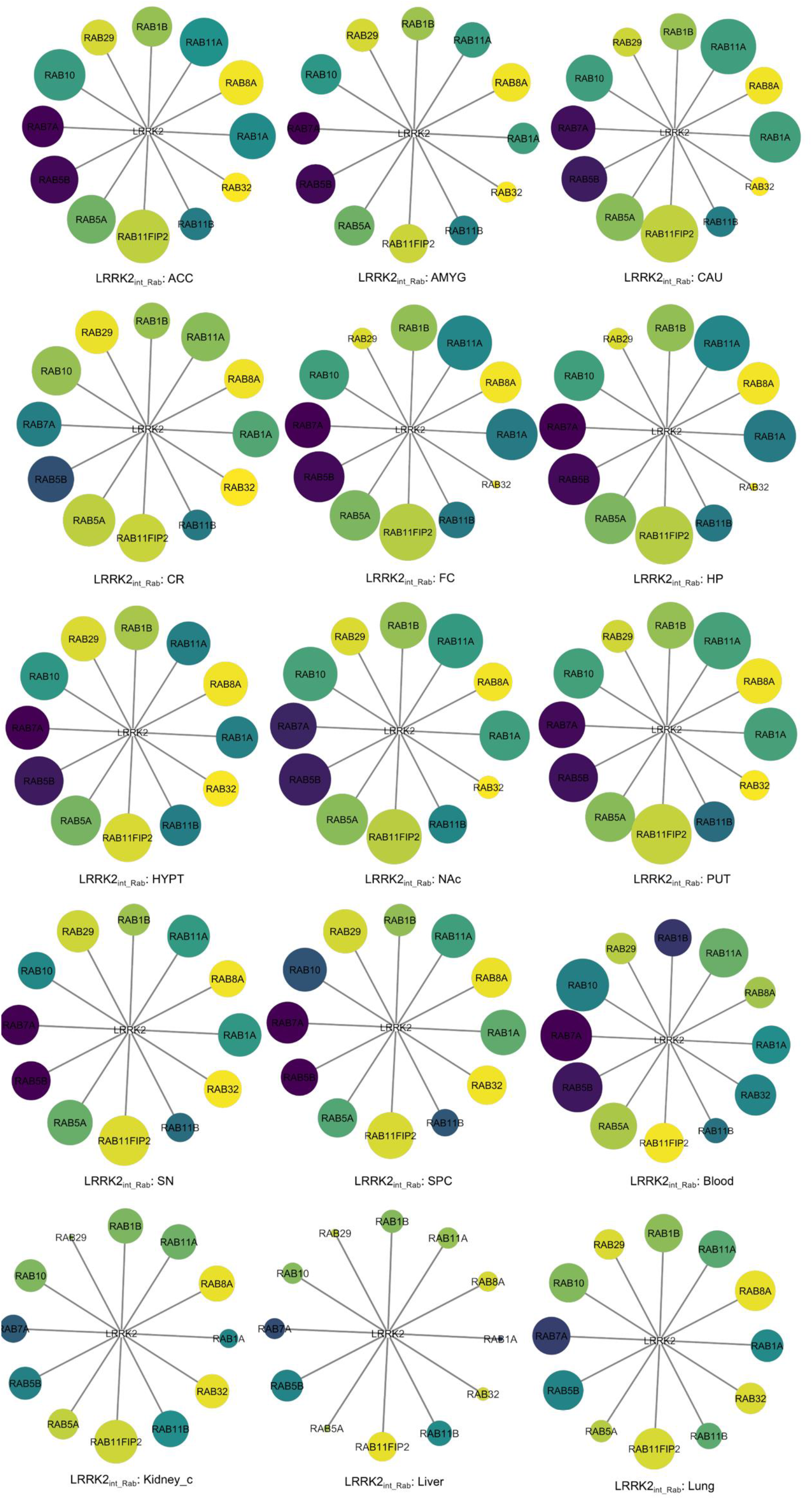
Tissue specific LRRK2:Rab interactomes (LRRK2_ints_Rab_). The network graphs show the different interaction attributes between LRRK2 and 12 Rab proteins derived from the general LRRK2_int_. Nodes represent Rab interactors of LRRK2. Node colour represents normalised expression levels of Rab proteins. The darker the colour, the higher the expression level of a certain Rab interactor of LRRK2 in a given tissue. Node size represents co-expression coefficients. The larger the node, the higher the co-expression coefficient presented between LRRK2 and a certain Rab interactor in a given tissue. Abbreviations: ACC: Anterior Cingulate Cortex; AMYG: Amygdala; CAU: caudate; CR: cerebellum; FC: frontal cortex; HP: hippocampus; HYPT: hypothalamus; NAc: nucleus accumbens; PUT: putamen; SN: substantia nigra; SPC: spinal cord c-1; Kidney_c: kidney cortex.

## Discussion

In this study, we constructed the LRRK2 protein-protein interactome from PPIs derived from peer-reviewed literature. To maximise literature coverage, PPIs were obtained from 3 secondary repositories: HIPPIE, MIST and PINOT. These tools extract PPIs from multiple primary, manually curated databases allowing to effectively text mine the available PPI literature. Although PINOT, MIST and HIPPIE are usually considered as interchangeable resources, they collect PPI information in different manners: PINOT extracts human PPIs from 7 primary databases (bhf-ucl, BioGRID, InnateDB, IntAct, MBInfo, MINT and UniProt) via the PSIQUIC interface, while in HIPPIE and MIST PPIs are firstly retrieved from several primary databases (including IntAct, MINT, BioGRID, HPRD, DIP, BIND and MIPS) and then stored in an internal repository. Although this strategy allows for a quicker retrieval of PPIs from HIPPIE and MIST in comparison to PINOT, their internal repositories require constant updates, while PINOT does not due to the live download via PSIQUIC. Another point of difference is in the quality control process these 3 tools apply to the PPI data. These processes involve divergent scoring systems based on different factors, thereby causing potential variation in the query results. Therefore, a combined use of HIPPIE, MIST and PINOT was essential to effectively maximise the literature coverage while extracting LRRK2 PPIs. A total of 3831 PPI annotations were collectively retrieved for LRRK2. After removal of duplicated entries, a list of 1871 unique interactions remained, suggesting indeed a high (but not complete) overlap in performance among the 3 tools used. Considering the total list of 1871 PPIs, 529 (28.3%) were retrieved from all 3 databases; 902 (48.2%) from 2 of the 3 tools; while 440 (23.5%) were obtained from one tool only (373 from MIST and 15 from HIPPIE)

With such a large number of LRRK2 interactors, it was important to set filters to extract the most reliable core within the vast LRRK2 interactome. We defined “reliability” as “reproducibility”, therefore we reduced the LRRK2 PPIs to those that have been reported at least twice in literature. To achieve this, we followed a 2-step strategy. Firstly, we converted similar interaction detection methods into one single method class (using the PINOT Method Grouping Dictionary, Table S1). This step was essential for accurate evaluations of the reproducibility of the LRRK2 PPIs, as curators behind different databases might record the same PPI under multiple interaction detection methods that are actually methodological synonyms, thereby introducing semantic bias. Secondly, “Low quality” PPIs – defined as interactions reported in literature within 1 publication only and validated with methods belonging to 1 category only – were filtered out from the LRRK2 interactome.

Following the 2-step strategy, the LRRK2 interactome was reduced to only 408 out of 1871 (21.8%), clearly showing that one of the problems we are facing with LRRK2 investigations (and potentially with PPI analyses in general) is that most of the data are not reproduced and thereby not directly trustable. This poor reproducibility may result from delays in curation of papers into primary databases, limited interest in wet-lab research aimed at reproducing information that has already been published, or the real inflation in type 1 errors within the field of protein interaction research.

The 408 interactors that passed the quality control constituted the general LRRK2 interactome, which is to our knowledge the most comprehensive, quality controlled human protein interactome of LRRK2 (LRRK2_int_) currently available. Further analysis of the general LRRK2 interactome was performed to evaluate whether the stringent, reductionist approach adopted in the construction of the general LRRK2 interactome was adequate to capture the current knowledge around LRRK2. Among the 408 interactors, LRRK2 itself exhibited the highest score for interaction (FS = 50), confirming LRRK2 self-interaction as the best known and reproducible PPI of LRRK2. Heterologous interactors with high FS (i.e. replicated in many publications using a plethora of different methods) were, as expected: HSP90AA1 (FS=19); YWHAQ (FS=14), HSPA8 (FS=13), MSN (FS=13), YWHAZ (FS=13), CDC37 (FS=11), DNM1L (FS=11), STUB1 (FS=11) and TUBB (FS=11). A large proportion of the general LRRK2_int_ was composed of cytoskeletal proteins (12.5%), which is in accordance with the known role of LRRK2 in cytoskeletal dynamics (reviewed by (Civiero *et al*, 2018)). Finally, 35 protein kinases and 13 Rab GTPases were part of the LRRK2_int_, confirming, again, that the general LRRK2_int_ built in our analysis does recapitulate the general knowledge related to LRRK2 interaction partners.

Functional enrichment analysis of the general LRRK2_int_ identified a plethora of biological processes, which were grouped into 14 larger functional groups. MCA was performed to compare the contribution of each single LRRK2 interactor to the enrichment of these functional groups. Interestingly, the majority of the functional groups (namely “autophagy”, “cell death”, “development”, “intracellular organisation”, “metabolism”, “transport” and “regulation of gene expression”) were not combined in any cluster. Only 2 clusters were identified: 1) the super-functional group of “protein metabolism” (Cluster I) containing “protein modification”, “protein localisation” and “protein catabolism”, and 2) the super-functional group of “response to stimulus” (Cluster II) comprising “signalling”, “regulation of enzyme functions”, “immune system” and “response to stress”. These results confirmed that i) the LRRK2_int_ is indeed involved in a wide range of different functions and ii) when looking at each of these functions, it appears that they are driven by distinct sets of LRRK2 interactors. In other words, LRRK2 interactors do not contribute, as a group, to the enrichment of all the LRRK2 associated functions. Rather, specific sets of LRRK2 interactors contribute to the enrichment of specific LRRK2 associated functions. This note is of importance as it sustains the hypothesis that differential expression of groups of LRRK2 interactors in different tissues might indeed be responsible for the specialisation of LRRK2 functions. In this model, the multiple functions the LRRK2_int_ is associated with are not all active at the same time/place; rather they are differentially relevant within different tissues, cell types and moments in time.

RNA-seq read counts for the LRRK2 interactors were downloaded from GTEx for 11 brain regions and 4 peripheral tissues. These data were used to differentiate the general LRRK2_int_ into 15 tissue specific LRRK2 interactomes. Two different strategies were applied based on pair-wise DEA (considering absolute expression levels) and pair-wise co-expression analysis (considering LRRK2:interactor co-expression levels). In general, the level of co-expression is regarded as a preferential measure as it is assumed that co-expressed proteins are co-regulated at a gene level and involved in the same biological processes. However, there is no reason to rule out an analysis based on DEA as an increased level of expression for a certain protein in a tissue might indicate increased demand for the biological processes in which that protein participate. Therefore, DEA and co-expression analysis were used to analyse 15 tissue specific LRRK2_ints_ following the assumption that LRRK2 interactors can increase or decrease in concentration in a tissue specific fashion following the unique functional requirements of each tissue. Similarly, in different tissues, LRRK2 can be co-expressed with different interactors depending on the functional unit that need to be activated to sustain the tissue specific functions.

We then used these results to compare the 15 tissues against each other thus effectively evaluating the different expression behaviours of the LRRK2_ints_ across different tissues. Firstly, we evaluated the hierarchical clustering of the 15 tissues to compare the expression behaviour of the LRRK2_ints_ in the 4 peripheral tissues vs the 11 brain regions. The results showed a clear difference in expression profiles (peripheral tissues vs CNS). In the DNA analysis the 4 peripheral tissues formed a distinct cluster meaning the LRRK2 interactors are in general expressed at different levels in the periphery in comparison with the CNS. In the co-expression analysis, the different peripheral tissues did not cluster together, however, they did not cluster with the brain regions either. In particular, when the brain regions were all already grouped into 3 clusters, the peripheral tissues were still left unclustered. This might suggest that from a co-expression perspective LRRK2 interactors behave similarly in some brain regions, while they show peculiar and unique behaviours in the 4 peripheral tissues.

Interestingly, the hierarchical clustering of the 11 brain regions based on the results from both DEA and co-expression analysis showed a similar result: putamen, caudate and nucleus accumbens were always found within the same cluster, suggesting the expression behaviour of the LRRK2int is very similar within these 3 regions both in terms of absolute expression levels of LRRK2 partners and LRRK2:interactors co-expression. This suggests the 3 regions might form a LRRK2 functional unit within the brain where the LRRK2_int_ sustains similar processes/functions.

Of note, putamen, caudate and nucleus accumbens form the striatum, target for the projection of dopaminergic neurons and one of the most affected brain regions during Parkinson’s disease (PD) progression. In fact, multiple studies have associated the degeneration of putamen and caudate with motor and non-motor PD symptoms (Wang *et al*, 2018; Playford *et al*, 1992; Manes *et al*, 2018); while nucleus accumbens, involved in mediating emotional and motivational processes such as rewarding experiences, impulsive and compulsive behaviours, might be implicated in the neuropsychiatric symptoms of PD (Hammes *et al*, 2019; Barbosa *et al*, 2019).

The results from the expression analyses were also used to compare and rank the LRRK2 interactors based on their expression behaviour across the 15 different tissues. A total of 124 interactors presented in the same differential expression cluster as LRRK2 (named DEA_Cluster_LRRK2_). This cluster represents the proteins within the LRRK2_int_ that share the same pattern of differential expression as LRRK2 in the majority of tissues under analysis. A smaller number of interactors (n = 77) were grouped in the Co-ex_Cluster_LRRK2_, which is composed of the proteins (within the LRRK2_int_) presenting with the highest co-expression level with LRRK2 in the majority of the analysed tissues. Of note, 30 interactors overlapped between these two clusters, meaning they showed both conserved co-expression with LRRK2 and similar expression profiles as LRRK2 across the majority of tissues. Due to this peculiar behaviour, these 30 interactors might be the gateway to understand the “constitutive” LRRK2 functions that are conserved in different districts. These 30 interactors mainly participate in the functions of “intracellular organisation”, “metabolism” and “regulation of gene expression”.

We then investigated the functional patterns of each tissue specific LRRK2_int_. Firstly, we defined “core interactors” within each tissue as LRRK2 interactors that either presented high expression levels or high co-expression behaviours with LRRK2 in that given tissue. The thresholds for defining “core interactors” were tissue specific and were set on normally-distributed expression levels and co-expression coefficients (obtained via transformation) to ensure a similar numbers of “core interactors” were identified for each tissue. The tissue-specific “core LRRK2 interactors” were functionally annotated. Results showed that in the substantia nigra, the brain region involved in the primary degeneration during PD progression (Zou *et al*, 2021), the core LRRK2 interactors were involved in the functions of “intracellular organisation”, “protein catabolism” and “transport”, potentially suggesting that among all the LRRK2 functions, the biological processes concerning the regulation of endocytosis and trafficking of vesicles to the lysosomes might be of greater importance at this brain location. When we looked at the core LRRK2 interactors for the “putamen-caudate-nucleus accumbens” LRRK2 functional unit we previously defined, we found they are principally involved in the functions of “metabolism”, “protein localisation”, “response to stress”, “regulation of gene expression” and “signalling”. Of note, the “core LRRK2 interactors” for lung were particularly involved in autophagy and immune functions. This finding is in accordance with previous studies indicating that LRRK2 may function in preventing alveolar type II (AT2) cells from adverse immune response caused by pulmonary fibrosis (Kuwahara & Iwatsubo, 2020).

However, it should be noted that the functional groups in our analysis were generated with only Gene Ontology Biological Processes types of terms and classified via an in-house grouping dictionary. Therefore, the functional annotations can be incomplete, which is unfortunately a general issue with functional enrichment approaches. Hence, these functional data should be interpreted as suggestive rather than definitive.

Finally, we complemented our analysis by investigating the Rab protein family as an example to show how the LRRK2_ints_ generated from our pipeline can be applied to answer a specific research question regarding a singular or a particular group of interactors of interest. We extracted information focused on the LRRK2 interactors belonging to the Rab protein family thus describing the LRRK2-Rab relationship (from the perspective of expression and co-expression behaviours) across different tissues. Rab proteins have been widely recognised as LRRK2 interactors. They have been reported to cooperate with LRRK2 in a number of cellular processes such as the regulation of endolysosomal functions, response to stress and vesicle trafficking (Bae & Lee, 2020; Eguchi *et al*, 2018). Here we used tissue specific LRRK2:Rab interactomes to describe how this cooperation between LRRK2 and Rab proteins may potentially vary across different tissues. Our results suggested that, in general, caudate, putamen and nucleus accumbens (that we previously defined as striatal LRRk2 functional unit) have the highest levels of LRRK2:Rab co-expression, hinting a potentially stronger association between LRRK2 and the Rab proteins in the striatum in comparison with other brain regions or peripheral tissues. Among these Rab proteins, RAB7A and RAB5B presented both high expression and high co-expression levels with LRRK2 in most of the tissues, suggesting these 2 Rab proteins may play a constitutive and conserved role in the LRRK2:Rab interactome across the whole body. In comparison, RAB32 and RAB29 presented unique LRRK2:co-expression patterns with higher co-expression levels in substantia nigra, hypothalamus, spinal cord c-1, cerebellum and blood, suggesting these 2 Rab proteins may be specific LRRK2 partners in these tissues only.

## Conclusion

PPIs can help in understanding the functional milieu around a hub protein, such as LRRK2 in this study. However, tissue specificity is generally not considered in these analyses. With this work we have designed a pipeline that makes use of expression data to provide indication of tissue specific differences within the interactome of a given hub protein (LRRK2 in this case). On one hand we provided evidence that brain tissues are different from peripheral tissues concerning expression patterns of the LRRK2_int_ and within the brain we defined a cluster composed of caudate, putamen and nucleus accumbens, where the LRRK2 interactors shows very conserved expression patterns. Additionally, we identified 30 LRRK2 interactors, showing the most conserved co-expression with LRRK2 and the most similar expression profile as LRRK2 across the majority of tissues analysed. We have also identified tissue specific LRRK2 interactors, as those few LRRK2 interactors showing particularly high expression levels in 1 or 2 tissues only. Finally, we have used this pipeline to differentiate the functional patterns of LRRK2_ints_ in different tissues. On the other hand, we presented an example with the Rab protein family, to illustrate how the LRRK2_ints_ generated via our pipeline (available for download) can be filtered for answering research questions regarding the tissue specificity of LRRK2 PPIs. As LRRK2 is a crucial target for PD treatment, with several small molecules currently in clinical trials, a better understanding of LRRK2’s tissue specific functionality has indeed become a research priority. This pipeline intends to be a rigorous bioinformatical attempt to raise awareness of the complexity and variability of LRRK2’s interplay with its interactors.

